## Supplemental Figs for "Linking individual differences in human primary visual cortex to contrast sensitivity around the visual field"

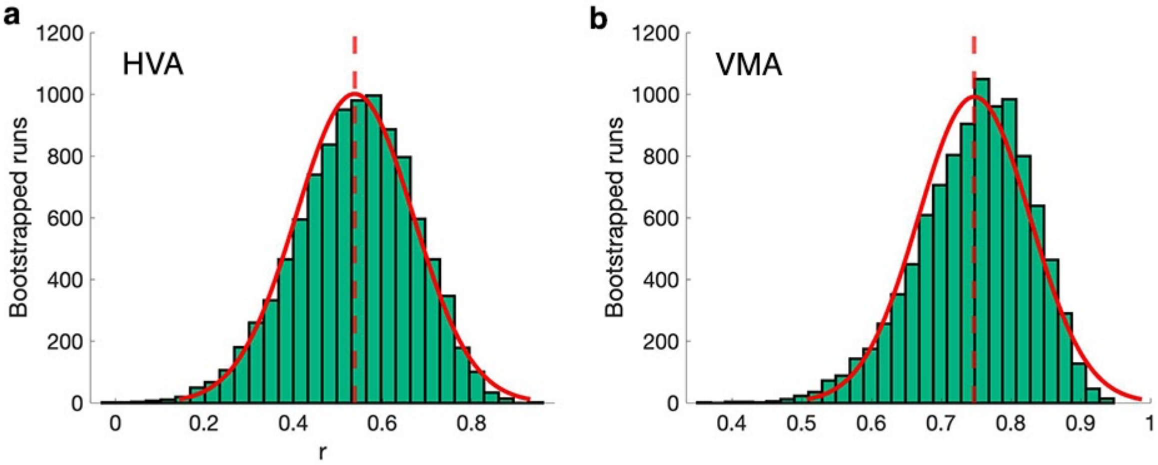


**Supplementary Fig 1.** Histograms of 10,000 bootstrapped Pearson's correlations between the **(a)** HVA and (**b)** VMA generated using subsampled blocks of contrast sensitivity data for each observer. For each observer, we randomly subsample and average two blocks of contrast sensitivity data and use this to generate HVA indices. The blocks are shuffled and this process repeated. We then correlate the pairs of HVA indices for each observer. This is repeated 10,000 times to generate a distribution of *r* values. The same is repeated for the VMA. The median *r* for the HVA is .55, and the median *r* for the VMA is .75, as denoted by the dashed red vertical line. The red line shows a normal distribution curve fit to the data. Source data for **(a)** and **(b)** are provided as a Source Data file.

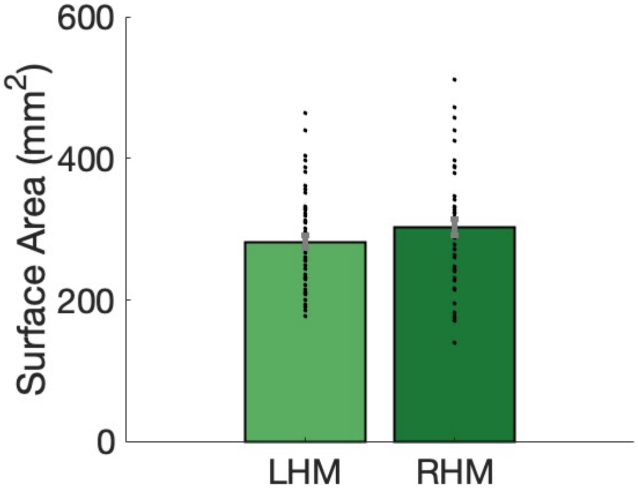


**Supplementary Fig 2.** The bias towards more surface area dedicated to processing ±15° along the right horizontal meridian when compared to the left horizontal meridian is marginally significant in an extended dataset (paired samples t-test, $t(53) = 1.89$, $p = .065$, $d = .26$). Data come from an extended dataset of 54 observers (29 included in the initial analysis and 25 additional observers). Error bars represent ±1 SEM and the gray error bar on the horizontal black bracket represents the standard error of the difference. Source data are provided as a Source Data file.

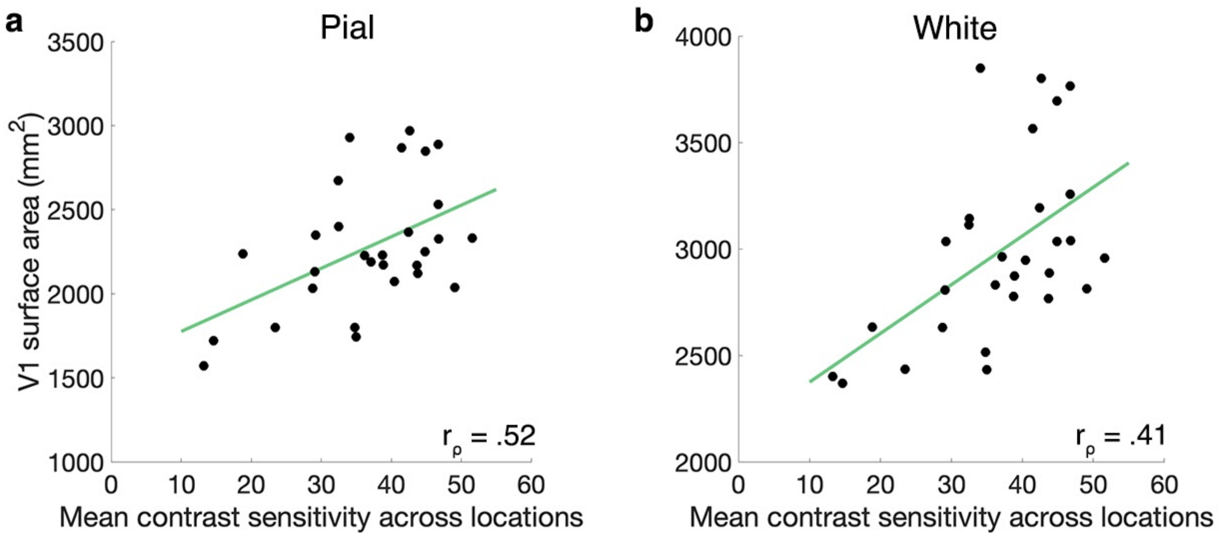


**Supplementary Fig 3.** Average contrast sensitivity across polar angle locations correlates with **(a)** V1 pial surface area (Spearman's correlation, one-tailed, $r_\rho = .52$, $p = .002$) and **(b)** V1 white matter surface area (Spearman's correlation, one-tailed, $r_\rho = .41$, $p = .014$) (n=29). Source data for **(a)** and **(b)** are provided as a Source Data file.

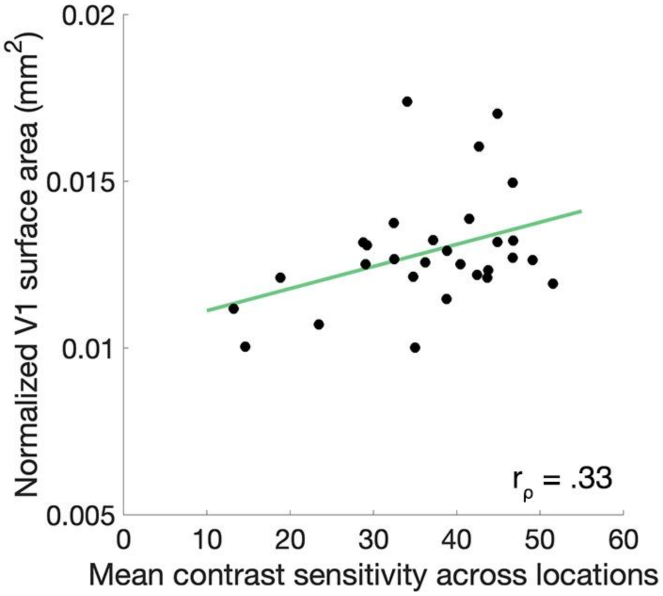


**Supplementary Fig 4.** The correlation between average contrast sensitivity across polar angle locations and V1 surface area is reduced after normalizing V1 surface area to cortical surface area (Spearman's correlation, one-tailed, $r_\rho = .33$, $p = .038$, n = 29). Source data are provided as a Source Data file.

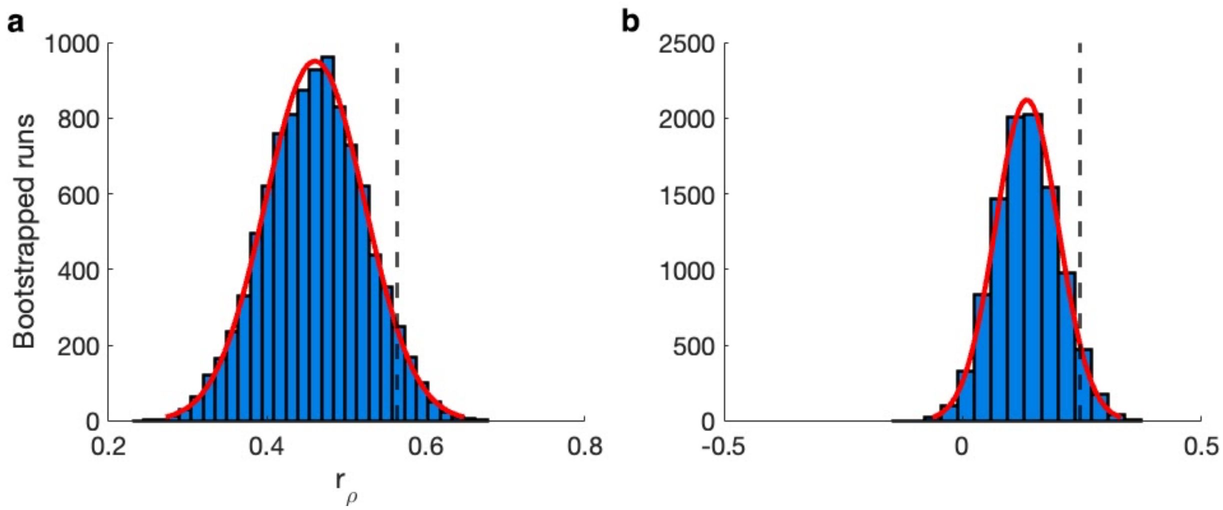


**Supplementary Fig 5.** Histograms of the null distribution generated by bootstrapping 10,000 Spearman's rho correlations. In **(a)**, the assignment of the four contrast sensitivity values and four local surface area values were shuffled across observers (with the tie between a given quadruple of contrast sensitivity or local surface area measurements maintained across locations). In **(b)**, the assignment of the four contrast sensitivity values and four local surface area values were shuffled across location (with the tie of a given quadruple of contrast sensitivity and local surface area measurements maintained across observers). The dashed black line indicates the $r_\rho$ value at the 95th percentile; for **(a)** $x_{0.95}$ = .56, and **(b)** $x_{0.95}$ = .24. The red line shows a normal distribution curve fit to the data. Source data for **(a)** and **(b)** are provided as a Source Data file.

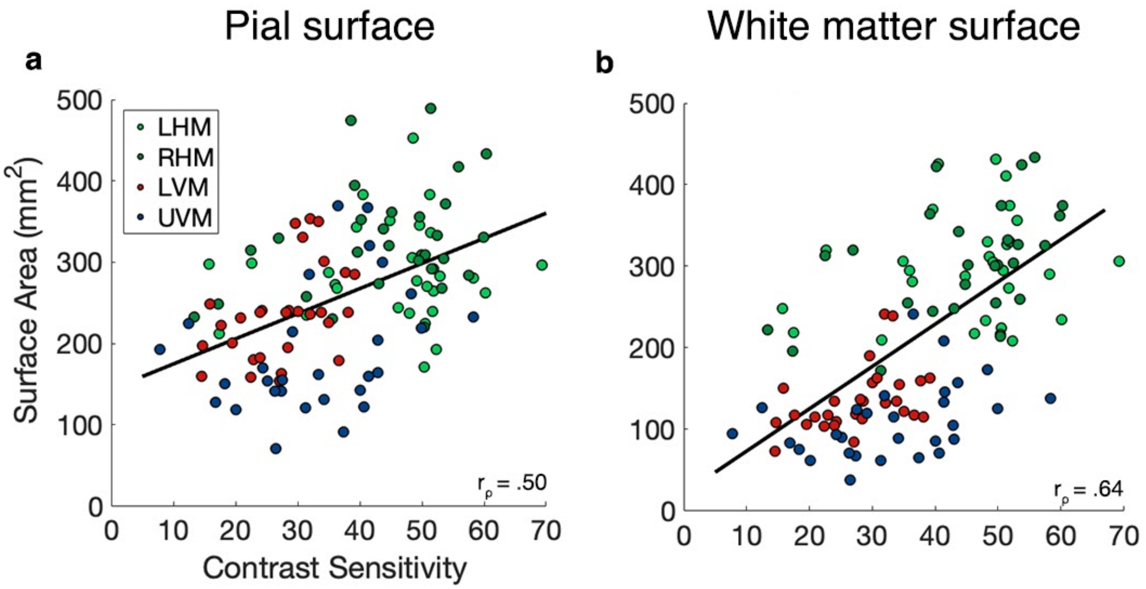


**Supplementary Fig 6.** Local contrast sensitivity correlates with local V1 cortical magnification measurements (calculated using the **(A)** pial (Spearman's correlation, one-tailed, $r_\rho$ = .50, $x_{0.95}$ = .46) and **(B)** white matter surfaces (Spearman's correlation, one-tailed, $r_\rho$ = .64, $x_{0.95}$ = .57) taken from the corresponding meridians (n=29). Source data for **(a)** and **(b)** are provided as a Source Data file.

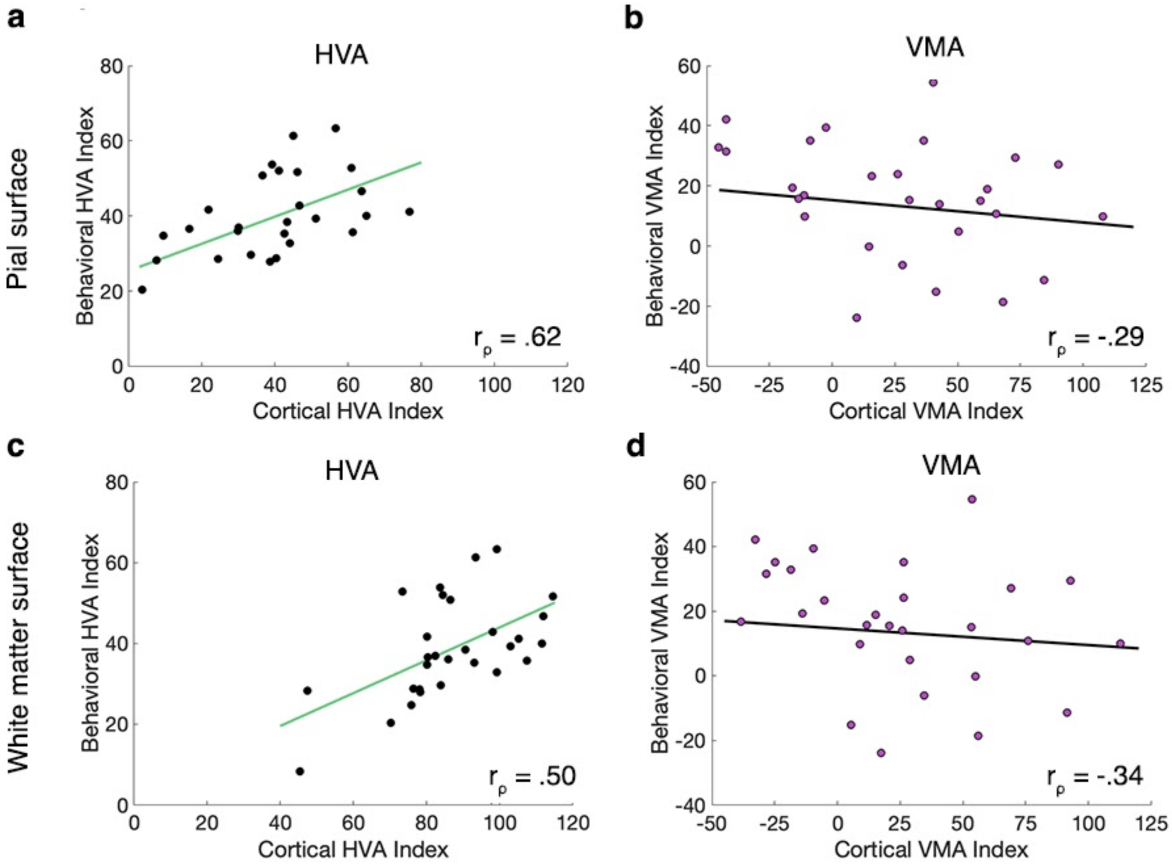


**Supplementary Fig 7.** Behavioral and cortical HVA and VMA correlations. In **(a)** and **(b)** the cortical HVA and VMA are calculated using the pial surface in **(c)** and **(d)** they are calculated on the white matter surface. The behavioral and cortical HVA (pial surface) correlate, (Spearman's correlation, one-tailed,$r_\rho$ = .62, $p <$ .001), as do the behavioral and cortical HVA (white matter surface) (Spearman's correlation, one-tailed,$r_\rho$ = .50, $p$ = .003). The behavioral and cortical VMA (pial surface) do not correlate (Spearman's correlation, one-tailed,$r_\rho$ = -.29, $p$ = .933, nor do the behavioral and cortical VMA (white matter surface) (Spearman's correlation, one-tailed,$r_\rho$ = -.34, $p$ = .965). Source data for **(a)**, **(b)**, **(c)**, and **(d)** are provided as a Source Data file.

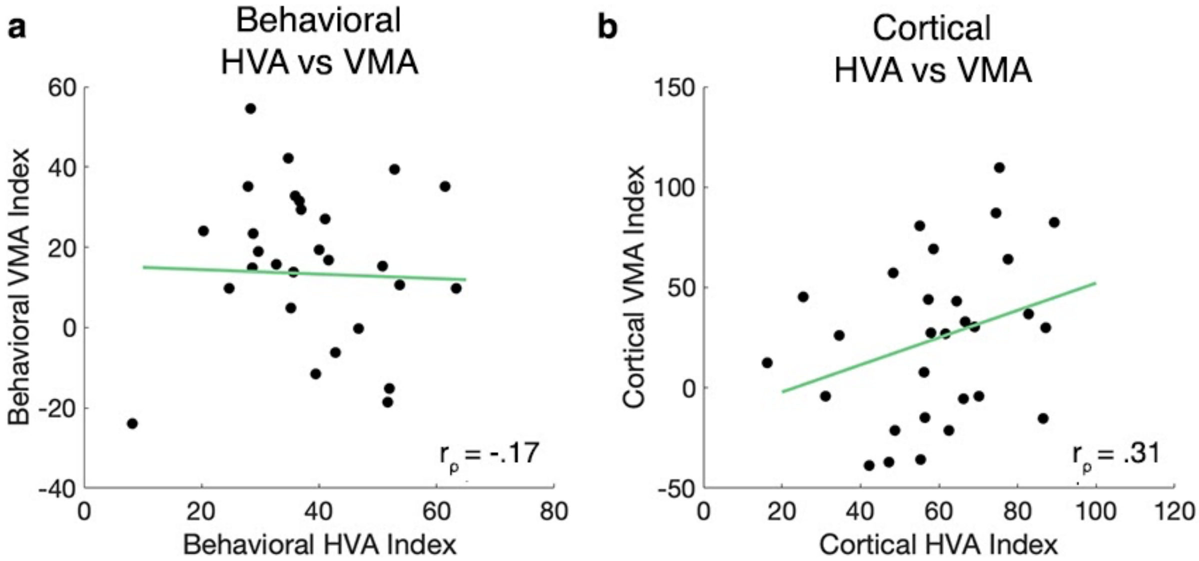


**Supplementary Fig 8. (a)** The behavioral HVA and VMA are not correlated (Spearman's correlation, one-tailed, $r_\rho$ = -.17, *p* = .810), **(b)** the cortical HVA and VMA (Spearman's correlation, one-tailed, $r_\rho$ = .31, *p* = .052) are marginally correlated. Source data for **(a)** and **(b)** are provided as a Source Data file.
